# A novel mouse model expressing human forms for complement receptors *CR1* and *CR2*

**DOI:** 10.1101/2019.12.23.887489

**Authors:** Harriet M. Jackson, Kate E. Foley, Rita O’Rourke, Timothy M. Stearns, Dina Fathalla, B Paul Morgan, Gareth R. Howell

**Affiliations:** The Jackson Laboratory, 600 Main Street, Bar Harbor, ME, USA; Dementia Research Institute Cardiff and Systems Immunity Research Institute, School of Medicine, Cardiff University, Cardiff, Wales, UK; Graduate School of Biomedical Sciences, Tufts University School of Medicine, Boston, MA USA; Graduate School of Biomedical Sciences and Engineering, University of Maine, Orono, ME, USA

**Keywords:** Complement cascade, complement regulators, immune cells, Alzheimer’s disease, Lupus

## Abstract

The complement cascade is increasingly implicated in development of a variety of diseases with strong immune contributions such as Alzheimer’s disease and Systemic Lupus Erythematosus. Mouse models have been used to determine function of central components of the complement cascade such as C1q and C3. However, species differences in their gene structures mean that mice do not adequately replicate human complement regulators, including *CR1* and *CR2*. Genetic variation in *CR1* and *CR2* have been implicated in modifying disease states but the mechanisms are not known. To decipher the roles of human *CR1* and *CR2* in health and disease, we engineered C57BL/6J (B6) mice to replace endogenous murine *Cr2* with human complement receptors, *CR1* and *CR2* (B6.*CR2CR1*). CR1 has an array of allotypes in human populations and using traditional recombination methods (*Flp-frt* and *Cre-loxP*) two of the most common alleles (referred to as *CR1*^*long*^ and *CR1*^*short*^) are replicated within this mouse model, along with a CR1 knockout allele (*CR1*^*KO*^). Transcriptional profiling of spleens and brains identifies genes and pathways differentially expressed between mice homozygous for either *CR1*^*long*^, *CR1*^*short*^ or *CR1*^*KO*^. Gene set enrichment analysis predicts hematopoietic cell number and cell infiltration are modulated by *CR1*^*long*^, but not *CR1*^*short*^ or *CR1*^*KO*^. Therefore, this mouse model provides a novel tool for determining the relationship between human-relevant *CR1* alleles and disease.

**Summary Statement:** We present the creation and validation of a novel mouse model that expresses human forms of complement cascade regulators CR1 and CR2.

## Introduction

The complement cascade is an integral component of our innate immune response and a first line of defense against bacterial infections. Various components of the complement cascade are constantly surveying for invading pathogens or debris, and tagging them for destruction. This system is composed of a number of plasma and membrane bound proteins and is tightly regulated. Circulating complement components are produced in the liver but can also be produced by specific cells in tissues. In recent years, the recognized roles of the complement cascade have expanded. For example, the complement cascade is integral for the process of synapse pruning during development and disease (Schafer and Stevens, 2010; Stevens et al., 2007), for regulation of embryo survival (Mao et al., 2003; Xu et al., 2000), and for tissue regeneration (Del Rio-Tsonis et al., 1998; Mastellos and Lambris, 2002; Rutkowski et al., 2010). Many of these novel roles were initially identified from animal models before being validated in human studies.

While animal models have proven fruitful in delivering greater understanding of the central components of the complement cascade such as C1q and C3, there are current limitations in studying human complement regulation in mice. In humans, the complement cascade is regulated in part by a series of genes on human chromosome 1 within the Regulators of Complement Activation (RCA) cluster (Hourcade et al., 1992; Lublin et al., 1987; Lublin et al., 1988; Rodriguez de Cordoba and Rubinstein, 1986; Rodriguez de Cordoba et al., 1985; Weis et al., 1987). A major difference in the RCA cluster between humans and mice is in the locus encoding Complement Receptor 1 (CR1/CD35), absent in mice (Farries and Atkinson, 1991; Jacobson and Weis, 2008; Nonaka, 2001) (**Fig. 1**). CR1 is both a receptor and a negative regulator of the complement cascade, binding to C3b, C4b, C1q, and MBL proteins. The interactions with C3b and C4b are considered to be the major function of this receptor (Iida et al., 1982; Klickstein et al., 1988; Krych-Goldberg et al., 1999; Wong et al., 1989; Zhang et al., 2013).

**Figure 1:**
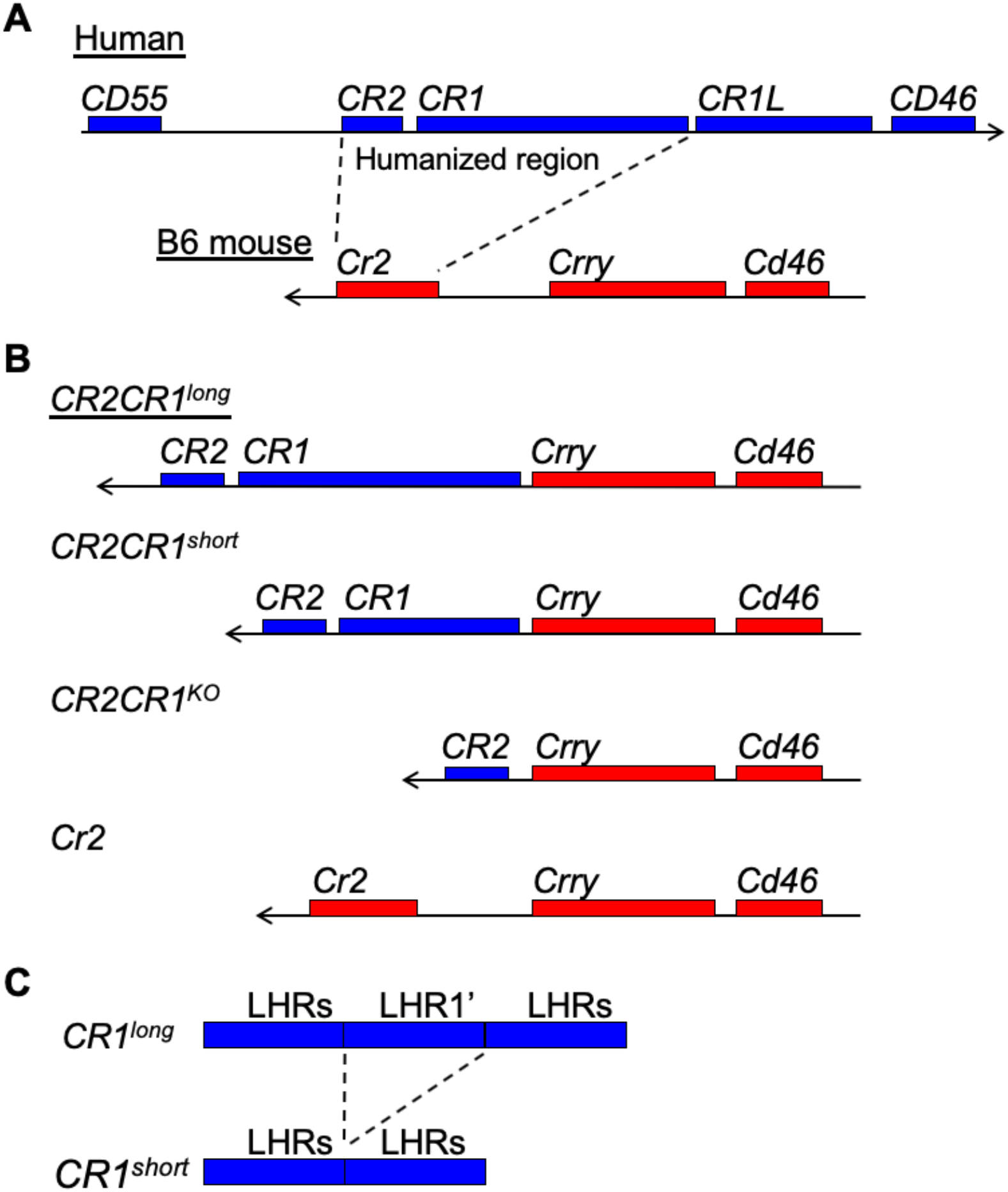
The B6. *CR2CR1* strain incorporates human *CR2* and common isoforms of human *CR1* into the mouse RCA cluster. (**A**) Humans and B6 mice differ in the genes they contain within the regulation of complement activation (RCA cluster). Human genes (blue boxes) include *CR2*, *CR1* and *CR1L* whereas B6 mouse genes (red boxes) include *Cr2* and *Crry*. (**B-C**) In the B6.*CR2CR1* strain, the mouse RCA cluster contains the human *CR2* gene and a long form of the human *CR1* gene (containing multiple Long Homologous Repeats, LHR). Through recombination using the *Flp/FRT* system, one LHR (LHR’) can be removed from the *CR1* gene. Through recombination using the *Cre/LoxP* system, human *CR1* can be deleted from the RCA cluster.

Genetic variation in *CR1* has been associated with a variety of diseases, such as Alzheimer’s disease (Biffi et al., 2010; Brouwers et al., 2012; Carrasquillo et al., 2010; Corneveaux et al., 2010; Hazrati et al., 2012; Jun et al., 2010; Keenan et al., 2012; Lambert et al., 2009; Van Cauwenberghe et al., 2013), Malaria (Birmingham et al., 2003; Khera and Das, 2009; Thomas et al., 2005; Xiang et al., 1999), and Systemic Lupus Erythematous (Birmingham et al., 2006; Corvetta et al., 1991; Holme et al., 1986; Katyal et al., 2003; Khera and Das, 2009; Marquart et al., 1995; Miyakawa et al., 1981; Moulds et al., 1996; Richardson et al., 1990; Ross et al., 1985; Walport et al., 1985; Wilson et al., 1986; Wu et al., 2007); however, in many cases the precise genetic variations have not been identified. CR1 has at least four allotypes: CR1-F, CR1-S, CR1-F’ and CR1-D (also known as CR1-A, CR1-B, CR1-C and CR1-D respectively) (Dykman et al., 1983a; Dykman et al., 1983b; Dykman et al., 1984; Dykman et al., 1985; Van Dyne et al., 1987). These four allotypes differ in size through loss or acquisition of long homologous repeats (LHRs; each comprising seven short consensus repeats [SCRs]); the commonest form, CR1-F (gene frequency 0.87 in Caucasians), comprises four LHR and a total of 30 SCRs while the CR1-S allotype (gene frequency 0.11) comprises five LHRs and a total of 37 SCRs (Holers et al., 1987; Wong et al., 1989). Some studies suggest different allotypes may be responsible for modifying risk for disease but currently there is no effective model system in which to test this (Biffi, 2012; Biffi et al., 2010; Bralten et al., 2011; Hazrati et al., 2012). To address this knowledge gap, we created a novel mouse model that enables the expression of different forms of human CR1 that we refer to as *CR1*^*long*^ (equivalent to CR1-S) and *CR1*^*short*^ (equivalent to CR1-F) (**Fig. 1**). While previous mouse models have relied on either the mouse *Cr2* or *Crry* genes or on transgenically expressed forms of CR1 to study the function of human CR1 in mice (Davoust et al., 1999; Killick et al., 2013; Maier et al., 2008; Manickam et al., 2010; Marchbank et al., 2002; Pappworth et al., 2012; Prodeus et al., 1998; Ramaglia et al., 2012; Repik et al., 2005; Wu et al., 2002), this is the first model to express *CR1* within the equivalent region of the mouse genome with human relevant promoter and regulatory sequences. Transcriptional profiling of the spleen and brain reveals significant differences in gene expression between mice carrying different allotypes of CR1 supporting the use of this new mouse model as a tool for studying CR1-dependent disease mechanisms.

## Results

### Chimeras Produce Viable, Construct-Carrying Pups with Successful Germline Transmission

To overcome species differences between mice and humans, we developed a new mouse model that, in the place of mouse *Cr2* (*mCr2*), expresses human *CR2* and *CR1* (**Figs. 1A-B** and **2**, see Methods). The B6.*CR2/CR1* mouse model is capable of expressing two isoforms of CR1 (*CR1*^*long*^ and *CR1*^*short*^, **Figs. 1B**) representing common allotypes predicted to be relevant to human disease (Biffi, 2012; Biffi et al., 2010; Hazrati et al., 2012). The difference between *CR1*^*long*^ and *CR1*^*short*^ is the number of LHR regions (**Figs. 1C**). To maximize relevance to human *CR1* regulation, we have incorporated the human intergenic region (HIR) between *CR2* and *CR1* gene (**Fig. 2**). To create the B6.*CR2CR1* strain B6 ES cells were targeted with a synthetic construct using recombineering (**Fig. 2A**, see methods). Twenty-eight chimeras, derived from two chimeric lines (5H4 and 5E2, **Fig. 2B**), were assayed for the presence of the construct. Chimeric mice carrying the synthetic construct were bred to B6^Tyr^, and all black progeny were genotyped to confirm transmission (**Fig. 2C**). No sex bias was seen with regards to transmission of the construct. Once germline transmission, and no sex bias, was confirmed the development of the *CR1* allelic series (*CR1*^*long*^, *CR1*^*short*^, *CR1*^−^) was performed (**Fig. 2C**). All genotypes developed through the allelic series were successfully bred to homozygosity through brother/sister mating. Allele-specific genotyping was used to determine the presence or absence of specific regions that defined each strain (m*Cr2*, *CR2*, *CR1*^*long*^, *CR1*^*short*^, *CR1*^*KO*^, HIR; **Fig. 3**, see methods). Of note, the initial establishment of the B6.*CR1*^*KO/KO*^ line proved difficult, as low numbers of homozygous mice were generated. However, once a male and female B6.*CR1*^*KO/KO*^ were identified, a mating pair was established, and the litter sizes were comparable to those of the other strains.

**Figure 2:**
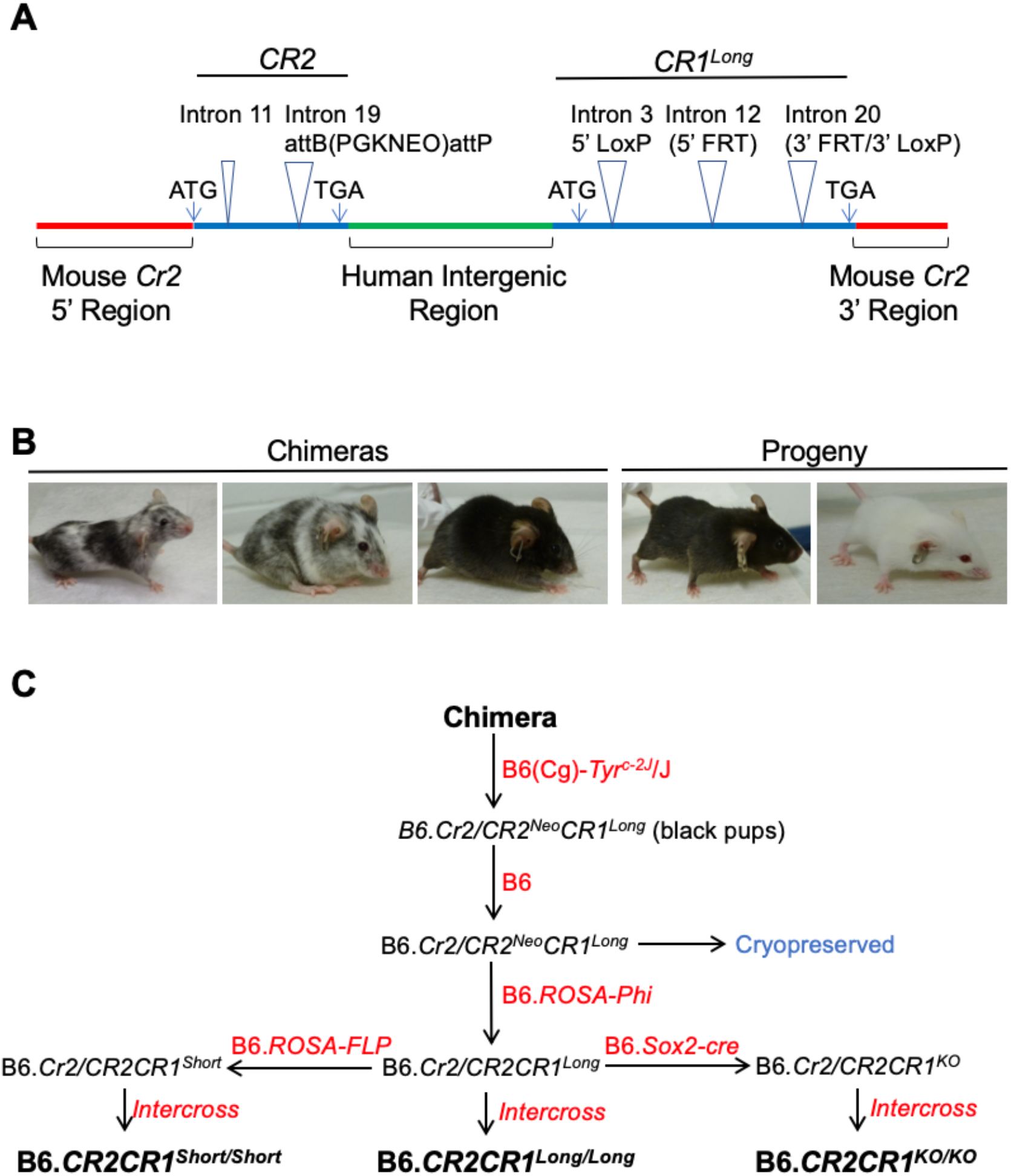
Creation of the B6. *CR2CR1* mouse strain. (**A**) The synthetic construct inserted into the B6 mouse genome (red bars) at the *Cr2* locus encompasses both the human *CR2* and *CR1* genes (blue bars), and included the human intergenic region between *CR2* and *CR1* (green bar). Human *CR2* sequence was based on NCBI reference sequence NM_001006658. Two introns (equivalent to introns 11 and 19) were included in human *CR2* gene. Intron 19 contained a neomycin cassette (PGKNEO) flanked by *AttB* and *AttP* sites. Human *CR1* sequence was based on NCBI reference sequence NM_000651. Three introns (equivalent to introns 3, 12 and 20) were included in the human *CR1* gene. *LoxP* sites were added to introns 3 and 20, *FRT* sites were added to introns 12 and 20. See *Supplemental File CR1 and CR2 protein alignments* for comparison of CR1^long^, CR1^short^ and CR2 protein sequences to reference protein sequence. (**B**) Example images of chimeras and progeny from the 5H4 ES cell line. Black pups were genotyped for the presence of the synthetic construct and used to establish subsequent strains (**C**) The breeding schemes to generate strains and experimental cohorts.

**Figure 3:**
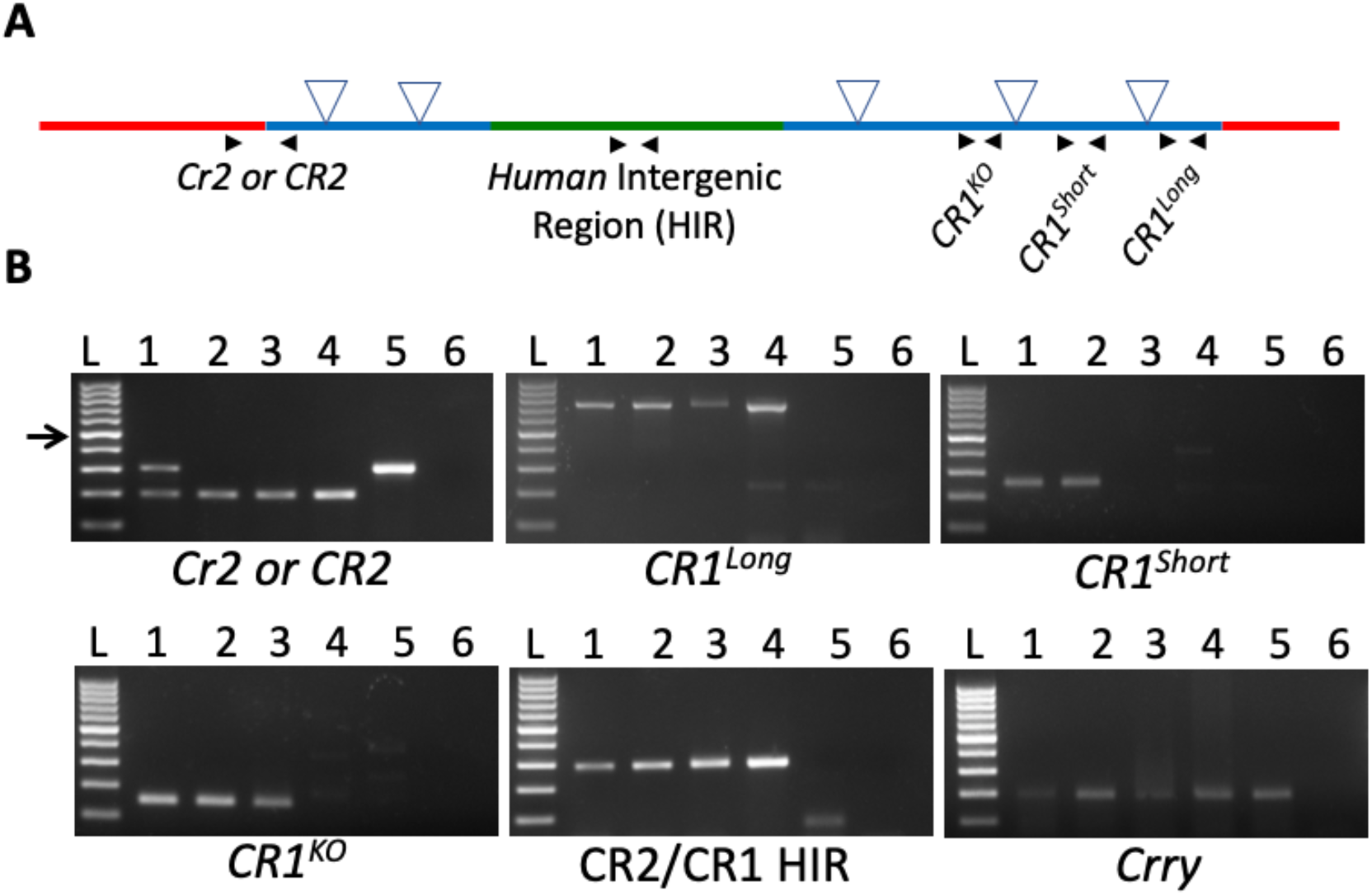
Validation of genomic regions within the B6. *CR2CR1* strains. (**A**) Location of primer pairs (black arrows) used to validate genomic regions. (B) The presence and absence of bands confirmed the presence or absence of each of the genomic regions and confirmed the *Crry* sequence has not been disrupted. Mouse *Cr2* = 300 bp product. Human *CR2* = 198 bp product. *CR1*^*long*^ = ~800 bp product. *CR1*^*short*^ (deletion between intron 12 and intron 20) = no product. *CR1*^*KO*^ (deletion between intron 3 to intron 20, **Fig. 2A**) = no product. *CR2/CR1* Human Intergenic Region (HIR) = 291 bp product. Crry = 190 bp product. L – Ladder (100 bp ladder, arrow is 500 bp). 1 – heterozygous B6.*CR2/CR1*^*long/+*^ with Neomycin cassette. 2 – homozygous B6.*CR2/CR1*^*long/long*^. 3 – homozygous B6.*CR2/CR1*^*short/short*^. 4 – homozygous B6.*CR2/CR1*^*KO/KO*^. 5 – B6. 6 – Water.

### RNA and protein expression of *CR1* and *CR2* observed in spleen of B6.*CR2CR1* mice

To establish RNA and protein expression of *CR1* and *CR2*, splenic tissue was assessed. As a primary organ of murine *Cr2* expression and complement-dependent immune complex processing, the spleen was an ideal target to validate the expression of human *CR1* and *CR2*. cDNA was generated from whole spleens of 3 males and 3 females from each genotype. Targeted primers confirmed the presence of human *CR2* and *CR1* in the spleen and the absence of mouse *Cr2* (**Fig. 4A-E**). For *CR1*, primers designed for *CR1* targeted both exon 2 and a region spanning exons 4 and 5. This strategy enabled identification of B6.*CR2CR1*^*KO/KO*^ mice that produced a transcript containing only the first two exons and not exons 4 and 5. As expected, *Crry* transcript, a mouse-specific gene that lies downstream of *Cr2*, was seen in all samples (**Fig. 4E**).

**Figure 4:**
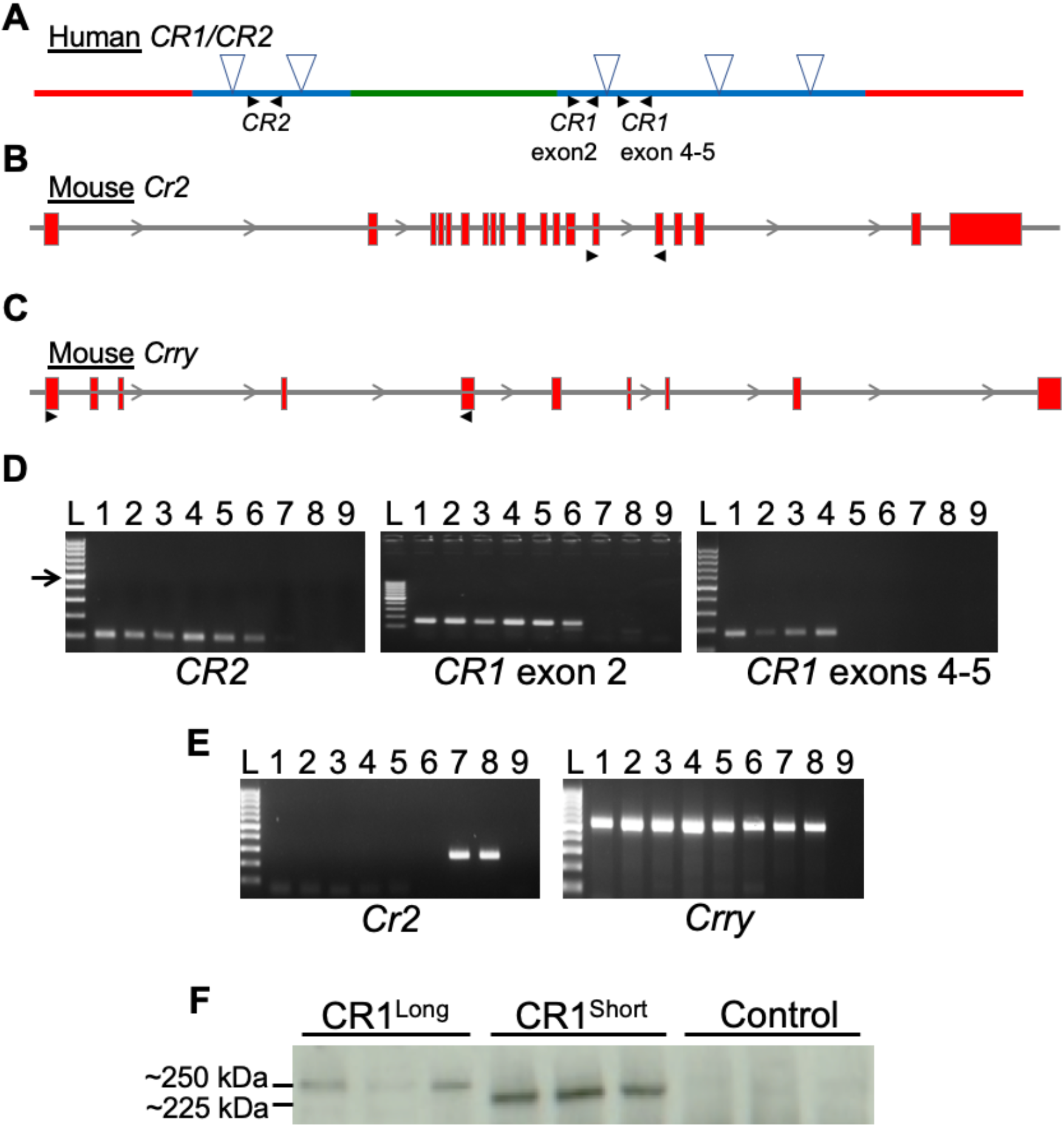
Generation of transcript and protein from the humanized region of the CR2CR1 mice in the spleen. (**A-C**) Each graphic depicts the humanized *CR2CR1* region (A), mouse *Cr2* gene (B) and mouse *Crry* gene (C). Black arrows denote the location of primer pair. (**D**) All B6.*CR2CR1* mice show expression of human *CR2* and *CR1* (exon 2). Only B6.*CR2/CR1*^*long/long*^ and B6.*CR2/CR1*^*short/short*^ show expression of *CR1* exons 4-5. (**E**) Only B6 mice show mouse *Cr2* expression. All mice show expression of mouse *Crry*. L-Ladder (100 bp ladder, arrow is 500 bp). 1 – Female B6.*CR2/CR1*^*long/long*^. 2 – Male B6.*CR2/CR1*^*long/long*^. 3 – Female B6.*CR2/CR1*^*short/short*^. 4 – Male B6.*CR2/CR1*^*short/short*^. 5 – Female B6.*CR2/CR1*^*KO/KO*^. 6 – Male B6.*CR2/CR1*^*KO/KO*^. 7– Female B6. 8 – Male B6. 9 - Water. Product sizes for RT-PCR are provided in the methods. (E) Western blot indicating mice expressing the humanized *CR2CR1* region produce protein products at their expected molecular weight. B6.*CR2/CR1*^*long/long*^ mice (CR1^long^) produce a product larger than that of their B6.*CR2/CR1*^*short/short*^ (CR1^short^) counterparts. No product was observed in B6 controls. Intensity of CR1 protein in B6.*CR2/CR1*^*short/short*^ mice appears greater than in B6.*CR2/CR1*^*long/long*^ mice.

To assess expression of human CR2 and CR1 protein isoforms, western blotting was used. Samples from B6.*CR2CR1*^*long/long*^ and B6.*CR2CR1*^*short/short*^ mice showed bands at their predicted sizes (273 kDa and 223 kDa respectively) (**Figs. 4F**, **S1** and **S2**). Interestingly, samples from B6.*CR2CR1*^*long/long*^ mice appear to have a lower level of protein expression compared to samples from B6.*CR2CR1*^*short/short*^ mice. Samples from B6 mice were used as a negative control and to determine cross reactivity of antibodies. For CR2, all mice showed a single band at 148 kDa, with the exception of B6 mice where a second 190kDa band was seen (**Fig. S2**). This is to be expected as the mouse *Cr2* gene produces two protein products via alternative splicing, in contrast to the human CR2 counterpart.

### CR1^long^ modifies expression of more genes in the brain and spleen compared to *CR1^short^*

Transcriptional profiling was performed to identify transcriptional differences between B6.*CR2CR1* and B6 mice. Spleen and brain samples from three male and three female B6.*CR2CR1*^*long/long*^, B6.*CR2CR1*^*short/short*^, B6.*CR2CR1*^*KO/KO*^ and B6 controls were assessed (24 samples in total). First, the spleen samples were analyzed (**Fig. 5A-C**). Compared to B6 controls, B6.*CR1*^*long/long*^ mice showed a greater number of differentially expressed (DE) genes than B6.*CR1*^*short/short*^ mice (104 and 38 genes respectively, **Tables S1 and S2**). Only ten genes were DE when comparing B6.*CR1*^*KO/KO*^ mice to B6 (**Table S3**) suggesting the human *CR2* gene functions similarly to mouse *Cr2*. Interestingly, the expression of the *CR1*^*short*^ transcript was almost twice as high as the *CR1*^*long*^ transcript (9.4 counts per million (cpm) compared to 5.6 cpm respectively) supporting our previous data that showed a greater amount of CR1^short^ protein in comparison to CR1^long^ protein (**Figs. 4F**, **S1** and **S2**).

**Figure 5:**
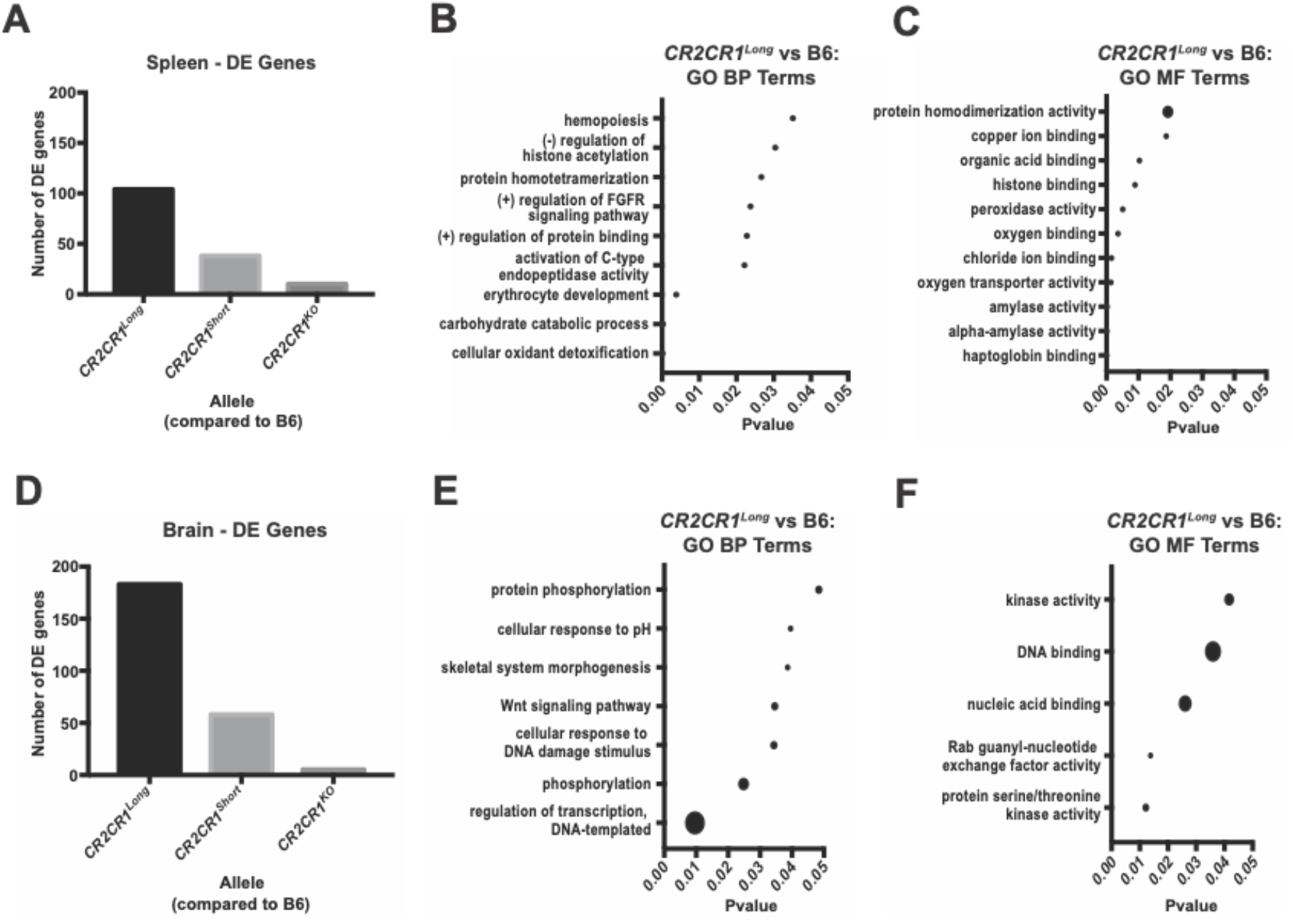
Transcriptional Profiling of the Spleen and Brain. (**A**) Number of differentially expressed (DE) genes (p<0.05) in the spleen for i) B6.*CR2CR1*^*long/long*^ compared to B6, ii) B6.*CR2CR1*^*short/short*^ compared to B6, and iii) B6.*CR2CR1*^*KO/KO*^ compared to B6. (**B**) Biological Process (BP) GO terms associated with the 104 DE genes between B6.*CR2CR1*^*long/long*^ compared to B6 in the spleen. (**C**) Molecular Function (MF) GO terms associated with the 104 DE genes between *CR2CR1*^*long/long*^ compared to B6. (**D**) Number of DE genes between each previous comparison in the brain. (E) Biological Process (BP) GO terms associated with the 183 DE genes between *CR2CR1*^*long/long*^ compared to B6. (F) Molecular Function (MF) GO terms associated with the 183 DE genes between B6.*CR2CR1*^*long/long*^ compared to B6. Dot size correlates to number of genes in the pathway associated with each term (see **Table S7**).

To predict functional relevance of 104 DE genes comparing spleen samples from B6.*CR2CR1*^*long/long*^ compared to B6, Kyoto Encyclopedia of Genes and Genomes (KEGG) and Gene Ontology (GO) term enrichment analyses were performed. Four KEGG pathways were significantly enriched (p<0.05), with two of the four relevant to amylase related genes (starch and sucrose metabolism, carbohydrate digestion and absorption) and the other two pathways (African trypanosomiasis and malaria) driven by ‘heme’ related genes. Enriched biological processes included metabolic terms such as ‘cellular oxidant detoxification’ and ‘carbohydrate catabolic process’. Interestingly, ‘negative regulation of histone acetylation’ was significant, suggesting that expression of CR1 in B6.*CR2CR1*^*long/long*^ mice may affect some epigenetic signatures when compared to B6 controls. For molecular function (MF), GO terms showed changes relating to binding, such as ‘haptoglobin binding’, ‘chloride ion binding’, ‘organic acid binding’ and ‘copper ion binding’, indicating that the *CR1*^*long*^ gene may be playing a role in intracellular binding (**Fig. 5C**). None of these pathways or GO terms were enriched in the DE genes comparing samples from B6.*CR2CR1*^*short/short*^ mice or B6.*CR2CR1*^*KO/KO*^ to B6.

The number of DE genes was greater in the brain when compared to the spleen (**Fig. 5D**). There were 183 DE genes identified by comparing B6.*CR2CR1*^*long/long*^ mice to B6 (**Table S4**), 58 DE when comparing B6.*CR2CR1*^*short/short*^ with B6 (**Table S5**), and only 5 DE genes between B6.*CR1*^*KO/KO*^ and B6 (**Table S6**). This trend reflects the results seen in the spleen, indicating that expression of *CR1*^*long*^ in mice had the greatest effect on gene expression in the brain compared with mice expressing either *CR1*^*short*^ or *CR1*^*KO*^. *CR1*^*long*^ was expressed at a low level in the brain (0.5 cpm on average), while no transcripts were detected for *CR1*^*short*^.

To identify the biological relevance of the 183 genes DE in brains of B6.*CR1*^*long/long*^ compared to B6, KEGG and GO term gene set enrichment was performed. Only one KEGG pathway, ‘Fanconi anemia pathway’ was significant (p<0.05) in the DE gene set between B6.*CR1*^*long/long*^ and B6 controls. Enrichment of this pathway was driven by the genes ‘*Wdr48’*, ‘*Atr’*, and ‘*Rev1*’ which respond to DNA damage. GO BP terms associated with differences between B6.*CR1*^*long/long*^ and B6 controls identified ‘regulation of transcription, DNA-templated’, ‘phosphorylation’, and ‘cellular response to DNA damage stimulus’ indicating DNA-damage response genes may be affected by the long allele. GO MF term analysis identified terms involved in kinase activity and DNA binding, suggesting potential further involvement in DNA repair mechanisms. No pathways or GO terms were enriched in mice expressing either *CR1*^*short*^ or *CR1*^*KO*^ (compared to B6) further supporting a model whereby CR1^long^ protein causes more changes to gene expression levels compared to CR1^short^.

### *CR1*^*long*^, but not CR1^short^, is predicted to modulate hematopoietic cell quantity and cell infiltration

To predict the functional consequences of the genes modified in B6.*CR2CR1*^*long/long*^, B6.*CR2CR1*^*short/short*^ and B6.*CR2CR1*^*KO.KO*^ mice compared to B6, ‘Disease and Function Analysis’ was performed in IPA. This function predicts increases or decreases in downstream biological activities using the direction of change of the genes in each DE gene list. In the spleen, multiple functional terms were predicted to be increased with the DE genes comparing B6.*CR2CR1*^*long/long*^ with B6. These could be generally classified as being related to regulation of hematopoietic cell number and were predicted to be activated (**Fig. 6A**). Two terms considered significant were ‘quantity of erythroid precursor cells’ and ‘quantity of myeloid cells’. Interestingly, genes associated with these terms were generally downregulated, including *Hba-a1* (−39.74 fold), *Hba-a2* (−19.25 fold) and *Hbb-bs (−20.00 fold)* which all encode for hemoglobin subunits (**Fig. 6B,C**). These data suggest that there is an effect of the *CR1*^*long*^ allele on hematopoietic quantity in the spleen. These functional consequences were not associated with DE genes when comparing samples from either B6.*CR2CR1*^*short/short*^ or B6.*CR2CR1*^*KO/KO*^ with B6.

**Figure 6:**
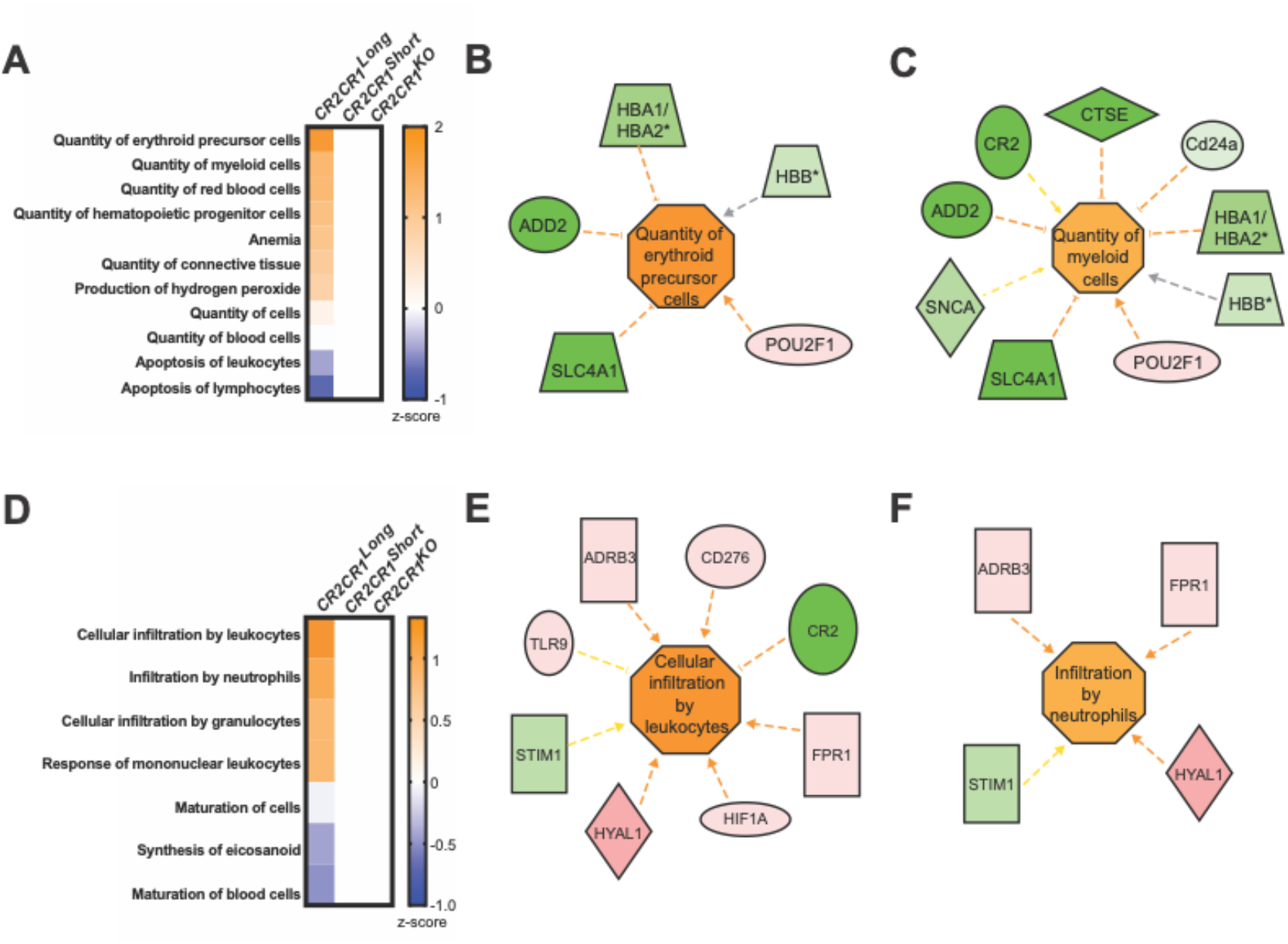
CR1^long^ is predicted to modulate hematopoietic cell quantity in the spleen and immune cell infiltration in the brain. (**A**) Disease and Function analytic (IPA) of transcriptional profiling data in the spleen identifies a significant positive association of DE genes in B6.*CR2CR1*^*long/long*^ compared to B6 samples with ‘Quantity of erythroid precursor cells’ and ‘Quantity of myeloid cells’ (p<0.05). (**B**) Genes associated with ‘Quantity of erythroid precursor cells’ associated genes and their effects. (**C**) Genes associated with ‘Quantity of myeloid cells’. (D) Disease and Function analytic (IPA) of transcriptional profiling in the brain identifies a significant positive association of DE genes in B6.*CR2CR1*^*long/long*^ compared to B6 samples with ‘Cellular infiltration by leukocytes’ and ‘Infiltration by neutrophils’. (E) Genes associated with ‘Cellular infiltration by leukocytes’. (F) Genes associated with ‘Infiltration by neutrophils’. Figures reproduced from IPA; shapes, colors, and color intensity follow IPA legend details. Orange = predicted activation, blue is deactivation. Red = increased expression in B6.*CR2CR1*^*long/long*^ compared to B6 samples. Green = decreased expression in B6.*CR2CR1*^*long/long*^ compared to B6 samples.

In the brain samples, functional consequences associated with DE genes comparing B6.*CR2CR1*^*long/long*^ with B6 samples included ‘cellular infiltration by leukocytes’ and ‘infiltration by neutrophils’ (p<0.05, **Fig. 6D**). DE genes associated with these terms were generally upregulated including *Hyal1* (14.78 fold), *Tlr9* (3.44 fold) and *Cd276* (4.59 fold) (**Fig. 6E,F**). HYAL1, a lysosomal hyaluronidase that degrades hyaluronan (a major constituent of extracellular matrix), has been shown to be involved in cell proliferation, migration and differentiation (Bharadwaj et al., 2007). TLR9, a toll-like receptor, plays a critical role in pathogen recognition and activation of innate immunity (Christensen et al., 2006; Santiago-Raber et al., 2009). CD276, also known as B7-H3, a member of the immunoglobulin superfamily, plays a role in T cell-mediated immune responses (Chapoval et al., 2001; Coyle and Gutierrez-Ramos, 2001). Collectively, these data predict CR1^long^ and CR1^short^ in the brain will differentially modulate immune-like cells such as resident microglia or the infiltration and functioning of peripherally derived immune cells – functional differences that may alter risk for disease such as Alzheimer’s disease.

## Discussion

Here, we present a new mouse model that may be of value to study of an array of human diseases including Alzheimer’s disease (Biffi et al., 2010; Brouwers et al., 2012; Carrasquillo et al., 2010; Corneveaux et al., 2010; Hazrati et al., 2012; Jun et al., 2010; Keenan et al., 2012; Lambert et al., 2009; Van Cauwenberghe et al., 2013), Systemic Lupus Erythematosus (Birmingham et al., 2006; Corvetta et al., 1991; Holme et al., 1986; Katyal et al., 2003; Khera and Das, 2009; Marquart et al., 1995; Miyakawa et al., 1981; Moulds et al., 1996; Richardson et al., 1990; Ross et al., 1985; Walport et al., 1985; Wilson et al., 1986; Wu et al., 2007) and infections such as malaria (Birmingham et al., 2003; Khera and Das, 2009; Thomas et al., 2005; Xiang et al., 1999). This model is the first to express both human proteins CR1 and CR2 in the mouse. We show that the humanized B6.*CR2CR1* strain expresses *CR2* in place of mouse *Cr2* and is capable of expressing two different isoforms of human *CR1* – a long form and a short form. CR1 and CR2 are important regulators of the complement cascade but their specific roles in human diseases have been difficult to study in models due to the species differences between humans and mice (**Fig. 1**) (Davoust et al., 1999; Killick et al., 2013; Maier et al., 2008; Manickam et al., 2010; Marchbank et al., 2002; Pappworth et al., 2012; Prodeus et al., 1998; Ramaglia et al., 2012; Repik et al., 2005; Wu et al., 2002).

We anticipate this mouse model will provide an important resource for elucidating the functions of CR1 and CR2 in human diseases – however further work is required to assess its full potential. First, it will be necessary to validate that human forms of CR1 and CR2 regulate the mouse complement system in a similar fashion to those seen in human studies. To do this, both CR2 and CR1 (long and short forms) will need to be tested for their ability to bind mouse C3b, C4b and C1q, three central components of the cascade. Second, testing existing and new antibodies that specifically recognize the human forms of CR1 and CR2 expressed in this mouse is required to confirm tissue and cellular distribution patterns in health and disease. Commonly used mouse anti-human CR1 monoclonal antibodies were tested but were unsatisfactory in the B6.*CR2CR1* mouse, likely because of technical issues in detecting mouse antibody binding in mouse tissues (data not provided). Some previous studies show available anti-CR1 and anti-CR2 antibodies are inconsistent between assays and tissues samples (Fonseca et al., 2016). Testing existing and new antibodies for human CR1 and CR2 in the B6.*CR2CR1* strain as well as in strains deficient in mouse CR2 protein (B6.*Cr2*^*KO*^) and mouse CRRY protein (B6.*Crry*^*KO*^) will eliminate the antibody specificity issue and potential for cross-reactivity due to similarities between these homologous genes. Establishing the gene expression patterns of human CR2 and CR1 in specific cell types, particularly in the spleen, brain and blood, will be necessary. Human CR1 is broadly expressed, albeit in varying quantities, on the plasma membranes of blood derived cells, including erythrocytes, eosinophil, monocytes/macrophages, B-lymphocytes, dendritic cells and a sub-set of CD4+ T-cells (Danielsson et al., 1994; Fang et al., 1998; Merle et al., 2015a; Merle et al., 2015b; Pascual et al., 1993; Rodgaard et al., 1991; Rodgaard et al., 1995; Weiss et al., 1989), on endothelia and numerous cell types in tissues. Erythrocyte CR1 plays an integral role in the clearance of soluble immune complexes, transporting them to macrophages in the spleen and Kupffer cells in the liver (Cosio et al., 1990; Craig et al., 2002), allowing for these cells to engulf and eliminate immune complexes (Skogh et al., 1985; van Es and Daha, 1984). The levels of CR1 expression on erythrocytes can differ due to a *HindIII* restriction fragment length polymorphism, which corresponds to a SNP in intron 27 of the *CR1* gene (Wilson et al., 1987). Interestingly, our preliminary analyses of transcript and protein expression of the long and short forms of CR1 suggest higher expression levels of the short form compared to the long form in the spleen (**Fig. 4**).

The B6.*CR2/CR1* mouse model provides an important platform for studying the function of single nucleotide polymorphisms (SNPs) that have been shown to modify CR1 and CR2 function and increase risk for human disease. Several exonic SNPs have been suggested to influence the stability of CR1 on erythrocytes, and thus mediate the high and low levels of expression (Xiang et al., 1999). While this variation is seen on erythrocytes, leukocyte expression does not show the same variability (Wilson et al., 1986). Given the current focus on developing treatments for Alzheimer’s disease, we expect the B6.*CR2/CR1* model to be a key resource to advance our understanding of how *CR1* risk alleles contribute to disease susceptibility. In 2009, a genome-wide association (GWA) study identified CR1 as a potential risk factor for Alzheimer’s disease (Lambert et al., 2009). This association was corroborated in 2010 (Carrasquillo et al., 2010), 2012 (Brouwers et al., 2012; Hazrati et al., 2012; Keenan et al., 2012) and 2013 (Van Cauwenberghe et al., 2013). The exact nature of the association of CR1 with Alzheimer’s disease is incompletely understood. One study (Hazrati et al., 2012) identified the risk for Alzheimer’s disease is most likely associated with the B allele of CR1 (CR1-B, equivalent to *CR1*^*long*^ in this study), with one copy of CR1-B carrying a 1.8x higher risk of disease over the CR1-A/A allele (equivalent to *CR1*^*short*^ in this study) and a faster rate of cognitive decline. These observations make determining the function of *CR1* specifically in the brain of key importance to fully elucidate its role in Alzheimer’s disease. Differences in neuronal morphology and distribution between CR1-A/A and CR1-A/B carriers were also reported, the former having a more filiform neuronal structure with CR1 expression that associated with the endoplasmic reticulum, whereas the latter had a more vesicular-like pattern of CR1 expression associating with lysosomes; reduced expression levels of CR1-B in comparison to that of CR1-A were also seen. Another study (Keenan et al., 2012) identified specific *CR1* SNPs (rs6656401 and rs4844609) that influenced rate of cognitive decline in Alzheimer’s disease in combination with *APOE* status. The latter SNP is associated with a single amino acid change in the C1q binding region of CR1; patients carrying both *APOE4* and rs4844609 showed a faster decline in episodic memory. While the functional implications of this coding SNP is yet to be determined, it may impact clearance of Aβ through interfering with C1q binding (Hazrati et al., 2012). Young adults who carry the *CR1* SNP rs6656401 had reduced grey matter volume in the entorhinal cortex (Bralten et al., 2011), an area associated with atrophy in AD patients (Braak and Braak, 1991; Hyman et al., 1984). Biffi *et al.* (2010) also saw drastic differences in entorhinal cortex volume of AD and MCI patients depending on their *CR1* genotype. The advent of precise gene editing in mice by methods such as CRISPR/CAS9 will allow these putative risk SNPs to be studied in the context of the human proteins.

Analysis of the transcriptional profiling data for the brain suggests that the CR1 isoforms differentially affect immune cell infiltration or immune cell activation, processes that have been shown to be important in disease susceptibility, onset and progression. Critically, the long form of CR1, CR1-B, the reported risk allele for Alzheimer’s disease, was associated with a DE gene signature indicating upregulation of pathways labelled ‘cellular infiltration by leukocytes’ and ‘infiltration by neutrophils’; we thus speculate that the association of CR1 long allele with Alzheimer’s disease might be explained at least in part, by altered immune cell infiltration into the brain. The analyses presented in this study was in young, healthy mice – further studies incorporating aging and Alzheimer’s disease pathologies, including amyloid and TAU, are required to further corroborate these findings.

In summary, the ability to more closely study *CR1* and *CR2* in a model system such as the mouse will facilitate our understanding of the role that these receptors, and the complement cascade more generally, play in a wide variety of diseases that show a strong immune component. A more complete understanding of the complement cascade, and its regulators, will lead to more targeted and personalized therapeutics for common diseases such as Alzheimer’s disease and lupus.

## Materials and Methods

### Mouse husbandry

All mice were maintained on a 12/12hr light/dark cycle. Mice were housed in 6-inch duplex wean cages with pine shavings, group-housed dependent on sex at wean, and maintained on LabDiet ^®^ 5K67. The Institutional Animal Care and Use Committee (IACUC) at The Jackson Laboratory (JAX) approved all mice used in this study. Daily monitoring of mice via routine health care checks was carried out to determine general wellbeing, with any mice considered to be unhealthy being euthanized with IACUC approved CO_2_ euthanasia methods.

### Humanizing Complement Receptors CR1 and CR2

The *CR2CR1* mouse model was created by Genetic Engineering Core at JAX via vector targeted embryonic stem (ES) cells. Due to the size, a multi-staged approach was used to create the targeting construct. Regions were designed *in silico* to encompass the human mRNA transcripts of *CR2* and *CR1* along with their corresponding human intergenic region (HIR). In parallel to this design a retrieval vector for mouse *Cr2* was utilized, targeted with a Spectinomycin (Spec) cassette, producing a vector with the 5’ and 3’ flanking regions of mouse *Cr2* (m*Cr2)*. To ensure the integrity of the human genes, they were assembled in a linear manner. The human *Cr2* mini gene was excised from its vector and incorporated within the HIR gap repair vector. This was then targeted to the m*Cr2*/Spec vector. The *CR1* mini gene containing vector was then targeted using *Apa1* and *AvrII* restriction enzymes, excising the fragment for integration into the multigene vector. Finally, this multigene vector was targeted with a Neomycin (Neo) cassette at synthetic intron 19 in the human *CR2* mini gene. Once the vector was confirmed, C57BL/6J (B6, Jax #664) ES cells were targeted, with incorporation occurring at the genomic locus for m*Cr2*. ES cells were transferred to a blastocyst from a B6(Cg)-*Tyrc-2J*/J (B6^Tyr^, Jax #58) and implanted into pseudo-pregnant females. Litters contained a variety of chimeric pups with differing degrees of penetrance.

Chimeric mice from two targeted B6 ES cell lines, 5H4 and 5E2, were bred to B6^Tyr^ mice (**Fig. 2**). From these, black pups were pursued for further breeding as a consequence of B6 ES cells being targeted initially. Mice determined positive through genotyping for the construct were then bred to B6 to confirm germline transmission by genotyping (**Fig. 2**). After germline transmission was confirmed, male mice from each line were sent for sperm cryopreservation, with the Neomycin (Neo) cassette intact, and are available upon request. Mice from the 5H4 line were used throughout this study.

### Developing the *CR1* allelic series

The initial stage of developing the allelic series was to remove the Neo cassette within intron 19 of *CR2*. To achieve this, mice derived from the 5H4 ES cell line were bred to B6.129S4-*Gt(ROSA)26Sortm3(phiC31*)Sor*/J (B6.ROSA-Phi, Jax #7743) mice, to target the attB-attP region surrounding Neo (**Fig. 2**). Mice negative for Neo were intercrossed to establish B6.*CR2CR1*^*long/long*^ strain (**Fig. 1 and 2**). For *CR1*^*short*^, B6.*CR2CR1*^*long/+*^ mice were crossed to B6.129S4-*Gt(ROSA)26Sortm1(FLP1)Dym*/RainJ (B6.ROSA-Flp, Jax #9086) mice. The Flp recombinase is ubiquitously expressed and targets the removal of LHR1’ (**Fig. 1**) encoded by exons 13-20 (via the flanking FRT sites in synthetic introns 12 and 20; **Fig. 2**). Mice carrying *CR1*^*short*^ (B6.*CR2CR1*^*short/+*^) were intercrossed to establish the B6.*CR2CR1*^*short/short*^ strain (**Figs. 1 and 2**). Finally, for *CR1*^*KO*^, B6.*CR2CR1*^*long/+*^ mice were crossed to B6.Cg-*Tg(Sox2-cre)1Amc*/J (*Sox2*-cre Jax #8454). In this strain, Cre recombinase is ubiquitously expressed and excises the targeted region using the LoxP sites, located within introns 3 and 20, to create a null allele (knockout, KO). Female B6.*Sox2*-cre mice were bred to male B6.*CR2CR1*^*long/+*^ mice, as Cre recombinase is active without necessarily needing to be inherited. Mice carrying *CR1*^*KO*^ (B6.*CR2CR1*^*KO/+*^) mice were intercrossed to establish the B6.*CR2CR1*^*KO/KO*^ strain (**Figs. 1 and 2**).

### Genotyping

PCR assays for genotyping were as follows:

For mouse *Cr2* or human *CR2* (**Fig. 3**): Common primer forward primer: 5’ - TCTTCCTCTCCTTGCTACAGG - 3’ C Cr2 Reverse: 5’ - AGAAGAGGTGGGGACGTTCT - 3’ and CR2 Reverse: 5’ - TACCAACAGCAATGGGGGTA - 3’ with the m*Cr2* product size at 300bp and the h*CR2* product size at 198bp with an annealing temperature of 60°C.

For the HIR Forward 5’ – TCACTCACCTCGAGCCATCT - 3’ and Reverse 5’ – TCAGCAGGTCTTGGCTTCAG – 3’ with a product size of 291bp at an annealing temperature of 59.3°C.

For *CR1*^*long*^ (**Fig. 3**): Forward 5’ – GTACTACGGGAGGCCATTCT – 3’ and Reverse 5’ – TGGCTTGGGGTACGCTC – 3’ with a product size of 708bp at an annealing temperature of 58.1°C.

For *CR1*^KO^ (**Fig. 3**): Forward 5’-TCTTGTACTACAGGGCACCG – 3’ and Reverse 5’ – ACCTCTAGGATTAAACGGTGGGG – 3’ with a product size of 150bp if cre recombination has not occurred, with an annealing temp 57.5°C. The absence of a band, with a CR1 positive genotype, indicates the removal of exons 4-20.

For *CR1*^*short*^ (**Fig. 3**), forward primer from the KO allele with the Reverse primer 5’ – CGATCATGGCTCACTGCGAA-3’. A product size of 251bp is expected if Flp recombination had not occurred. The absence of a band in a combination with a CR1 positive genotype, indicated Flp recombination. The annealing temperature for this reaction was 57.8°C.

For *Crry*: Forward 5’-TTGCTAATTGGTAGTGAGGAAAGG −3’ and Reverse 5’-TAAGTTGTTGTGAGGCTTGGGT −3’ with a product size of 190bp and an annealing temperature of 55.4°C.

### Cohort generation

Homozygous mice of the three genotypes *CR2CR1*^*long/long*^, *CR2CR1*^*short/short*^, CR2CR1^KO/KO^ were identified. A separate B6 colony was established for wild-type control samples. Cohorts of at least 4 males and 4 females were used for all assays except for transcriptional profiling where 3 males and 3 females per genotype were assessed. All mice were bled via submandibular bleed and tissue harvested at 3 months of age.

### Tissue harvesting and preparation

Mice were terminally anaesthetized using a Ketamine/Xylazine (99mg/kg Ketamine, 9mg/kg Xylazine) mix. They were transcardially perfused with PBS (phosphate buffered saline pH 7.4). Spleens and brains were harvested, snap frozen and stored at −80°C for further use. RNA and protein were extracted from the snap frozen tissue using Trizol according to manufacturer’s instructions. RNA was reconstituted in dH2O and protein was resuspended in 1:1 1% SDS/8M Urea. All RNA and protein samples were stored at −80°C before use. RNA concentrations were determined via Nanodrop and protein concentrations via DC Protein Assay respectively.

### cDNA synthesis and reverse transcriptase (RT)-PCR from RNA extracted from spleen

RNA extracted via Trizol was treated with DNase at 37°C for 30min, the reaction was stopped by placing on ice and 0.5M EDTA was used to deactivate the DNase. Samples were centrifuged and the supernatant was transferred to a new tube. A lithium chloride:ethanol solution was used to precipitate the RNA overnight at −20°C. Samples were centrifuged at maximum speed for 20min at +4°C, the supernatant removed and remaining pellets were washed with 70% ethanol. RNA was resuspended in dH_2_O and concentrations were read using the Nanodrop. 1μg of RNA was used to synthesize cDNA. Briefly, RNA was combined with random primers, dNTPs, RNase inhibitor, Multiscribe Reverse Transcriptase and made up to volume with dH_2_O. The reaction was incubated at 25°C for 10min, 37°C for 2hr, 85°C for 5min and +4°C. Samples were diluted 1:4 and concentrations were read again on the Nanodrop to ensure that no degradation had occurred. Samples were stored at −20°C until required. 100ng of cDNA was used to determine expression within the spleen. PCR assays for RT-PCR were as follows: For *CR2* at exon 11: Forward: 5’-TGGGGCAGAAGGACTCCAAT −3’ and Reverse: 5’-GCTCCACCATGGTCGTCATA −3’ with a product size of 148bp and an annealing temperature of 60°C.

For *CR1* at exon 2: Forward: 5’-TCCATTTGCCAGGCCTACCA −3’ and Reverse: 5’-TGCACCTGTCCTTAGCACCA −3’ with a product size of 152bp and an annealing temperature of 60°C.

For *CR1* spanning exons 4 and 5: Forward: 5’-TGGTTCCTCGTCTGCCACAT −3’ and Reverse: 5’-AGGATTGCAGCGGTAGGTCA −3’ with a product size of 178bp and an annealing temperature 60°C.

For m*Cr2*: Forward: 5’-TCATGAGGGTACCTGGAGTCA −3’ and Reverse: 5’-AAGAGGAATAGTTGACCGGTATTT −3’ with a product size of 244bp and an annealing temperature of 60°C.

For *Crry*: Forward: 5’-GGAGGAGTCAAGCTAGAAGTTT −3’ and Reverse: 5’-GTGTTGCAGCGGTAGGTAAC −3’ with a product size of 521bp and an annealing temperature of 55.3°C.

### Western Blotting

To determine protein presence and the size difference between the CR1^long^ and CR1^short^, alleles, 6% SDS PAGE gels were hand cast. Protein was diluted to 80μg of total protein with 2x Laemmli buffer (BioRad). Samples were denatured at 95°C for 5min and loaded onto the gel. Gels were run for 1hr at 150V and transferred to nitrocellulose membrane via the iBlot for 13mins. Blots were incubated at room temperature for 1 hour (hr) with blocking solution (5% skimmed milk powder block in 0.1% PBS-Tween), washed with 0.1% PBS-Tween for three 15min incubations and then incubated with rabbit-anti-human CR1 (also known as CD35; Abcam #ab126737, 1:100) for 48hrs in 0.1% PBS-Tween on an orbital shaker at +4°C. Blots were washed three times in 0.1% PBS-Tween and incubated with the appropriate secondary (Anti-Rabbit IgG HRP 1: 50,000) for 1.5hrs at RT. Blots were then washed an additional three times and detection was carried out using ECL detection regents (GE Healthcare). When required, blots were stripped by treatment with 0.25% sodium azide for 2hrs at RT and washed thoroughly in 0.1% PBS-Tween. Blots were re-blocked and re-probed with mouse anti-CD21 (anti-human CR2; Abcam #ab54253, 1:100) in 0.1% PBS-Tween overnight at +4°C. Blots were washed and incubated in the appropriate secondary antibody (Anti-Mouse IgG HRP 1:40,000), washed and detected. Finally, blots were treated with 0.25% sodium azide and probed with a loading control, anti-Vinculin (1:10,000) in 0.1% PBS-Tween overnight at +4°C, washed three times, incubated with the appropriate secondary antibody (Anti-Rabbit HRP 1:50,000) for 1hr at RT, washed and detected.

### Transcriptional profiling

#### RNA isolation, library preparation and sequencing

RNA was isolated from tissue using the MagMAX mirVana Total RNA Isolation Kit (ThermoFisher) and the KingFisher Flex purification system (ThermoFisher). Tissues were lysed and homogenized in TRIzol Reagent (ThermoFisher). After the addition of chloroform, the RNA-containing aqueous layer was removed for RNA isolation according to the manufacturer’s protocol, beginning with the RNA bead binding step. RNA concentration and quality were assessed using the Nanodrop (Thermo Scientific) and the RNA Total RNA Nano assay (Agilent Technologies). Libraries were prepared by the Genome Technologies core facility at The Jackson Laboratory using the KAPA mRNA HyperPrep Kit (KAPA Biosystems), according to the manufacturer’s instructions. Briefly, the protocol entails isolation of polyA-containing mRNA using oligo-dT magnetic beads, RNA fragmentation, first and second strand cDNA synthesis, ligation of Illumina-specific adapters containing a unique barcode sequence for each library, and PCR amplification. Libraries were checked for quality and concentration using the D5000 assay on the TapeStation (Agilent Technologies) and quantitative PCR (KAPA Biosystems), performed according to the manufacturers’ instructions. Libraries were pooled and sequenced by the Genome Technologies core facility at The Jackson Laboratory, 100 bp paired-end on the HiSeq 4000 (Illumina) using HiSeq 3000/4000 SBS Kit reagents (Illumina).

#### Sequence Alignment & Statistical Analysis Methods

FASTQ files were trimmed using Trimmomatic v0.33 which removed adapters and sequences with more than 2 mismatches, a quality score < 30 for PE palindrome reads, or a quality score of < 10 for a match between any adapter sequence against a read. A sliding window of 4 bases was used with a required (average) minimum quality score threshold of 15. The leading and trailing minimum quality score thresholds were set to 3 to keep a base. Reads had to be 36 bases or greater in length. Sequence alignment was completed using the *Mus musculus* Ensembl v82 reference genome in addition to three custom references based on that genome. Samples from both brain and spleen tissue were edited to include one (or none) of three human transgenes: *CR1*^*long*^, *CR1*^*short*^, or *CR1* KO. Each of the three human transgene sequences was appended to the *Mus musculus* reference genome individually; thus, three additional reference genomes were created to quantitate gene and isoform expression levels using RSEM v1.2.19. B6 samples were evaluated with the base *Mus musculus* reference genome. RSEM leveraged Bowtie2 alignment with strand-specific and paired-end parameters. The resulting gene/transcript (feature) count data were processed using edgeR v3.14.0 (R v3.3.1). For each of the four datasets (brain and spleen, gene and transcript), feature differential expression was evaluated in three pairwise comparisons per sex: 1.) *CR1*^*long*^ vs. B6, 2.) *CR1*^*short*^ vs. B6, and 3.) *CR1*^*KO*^ vs. B6. There were no differences between male vs female for each of the three comparisons, so samples were pooled by genotype to increase the quantity from three (three per sex) to six (pooled sexes). Any feature that did not have at least 1 read per million for at least 2 samples in either sets of samples evaluated in the pairwise comparison was excluded from the differential expression analysis. The Cox-Reid profile-adjusted likelihood method was used to derive tagwise dispersion estimates based on a trended dispersion estimate. The GLM likelihood ratio test was used to evaluate differential expression in pairwise comparisons between sample groups. The Benjamini and Hochberg’s algorithm was used to control the false discovery rate (FDR). Features with an FDR-adjusted p-value < 0.05 were declared statistically significant.

#### Gene set enrichment

The Database for Annotation, Visualization and Integrated Discovery (DAVID, v6.8) was used on each significant DE gene list for each pairwise comparison to identify enrichment of Kyoto Encyclopedia of Genes and Genomes (KEGG) pathways and Gene Ontology (GO) terms. Background gene sets were all trimmed normalized gene reads. KEGG pathways and GO terms were considered enriched with a p-value less than 0.05 (p<0.05). Ingenuity Pathway Analysis (IPA) was used for disease and functional analysis.

## Acknowledgements

The authors would like to thank The Jackson Laboratory Genetic Engineering Technologies for support in creating the B6.*CR2CR1* mouse model. They also thank Dr. Vivek Philip for assistance with analysis of transcriptional profiling data.

## Data Availability

All transcriptional profiling data is being submitted to Geo Archives and will be made available upon request.

## Author Contributions

HMJ, BPM and GRH conceived the study. HMJ performed experiments. ROR developed mouse cohorts. TMS and KEF performed analysis of transcriptional profiling data. DF validated and provided antibodies. HMJ, KEF, GRH and BPM wrote the manuscript. All authors approved the submission of the manuscript.

